# Computational model of primitive nervous system controlling chemotaxis in early multicellular heterotrophs

**DOI:** 10.1101/2024.08.30.610546

**Authors:** Egor O. Vasilenko, Anton V. Sinitskiy

**Affiliations:** RAS Institute of Molecular Biology, 119991, Moscow, Vavilov str., 32, Russia; ML LC, Massachusetts, USA

## Abstract

This paper presents a model to study a hypothetical role of a simple nervous systems in chemotaxis in early multicellular heterotrophs. The model views the organism as a network of motor units connected by flexible fibers and driven by realistic neuron excitation functions. Through numerical simulations, we identified the parameters that maximize the survival time of the modeled organism, focusing on its ability to efficiently locate and consume food. This synchronization enhances the ability of the modeled organism to navigate toward food and avoid harmful conditions. The model is described using basic mechanical principles and highlights the relationship between motor activity and energy balance. Our results suggest that even early prototypes of neural networks might provide significant survival advantages by optimizing movement and energy use. This study offers insights into how the first primitive nervous systems might have functioned. By publishing the code used in the simulations, we hope to contribute to the toolkit of computational methods and models used for exploration of neural origin and evolution.

## INTRODUCTION

The problem of the origin of the nervous system and the properties of its earliest evolutionarily examples remain an active area of research, with many questions still unanswered [1–10]. One possible scenario is the coordination of motion in multicellular organisms for more efficient locomotion, which, in its turn, may lead to more efficient feeding or predator evasion [6, 11]. Mathematical models, such as the Kuramoto model [12], have been applied to understand motor and sensory coordination in simple organisms with early nervous systems [13]. This model describes synchronization in coupled oscillators and has been applied to explain the coordinated movement of cilia and flagella in marine organisms, like Volvox colonies [14]. Discrete cellular automata models also simulate interactions in multicellular structures, such as Dictyostelia and Myxobacteria, helping to understand how synchronized movements can arise from local interactions without centralized control. The construction of decentralized elastic mobile structures is useful not only in biology, but also in robotics [15]. Research on jellyfish [16] reveals that their radially symmetric neural networks facilitate efficient movement and feeding. As the size increases during evolution, predators abandon temporal sampling in favor of spatial sampling, because the large length of the body contour in flat organisms or the surface area in volumetric organisms allows them to assess the gradient of food concentration [17]. However, the mathematical complexity of these systems presents challenges. While studies on more advanced organisms have advanced our understanding, the synchronization mechanisms in simpler life forms remain unclear. Early multicellular organisms, like choanocytes, likely developed neural precursors for coordination, but their evolutionary roles are underexplored. The ancient common ancestor of cnidarians and bilaterians had voltage-gated Na^+^ and Ca^2+^ channels [18]. Existing models often focus on specific aspects of synchronization and motor dynamics, leaving gaps in understanding the evolution of sensory-motor coordination across species. Modern species with nervous systems already have their differentiation, and, for example, the mechanism of action potential propagation has been investigated, but little is known about the first nervous systems and their emergence. This study aims to integrate synchronization dynamics, mechanical forces, and energy balance into a unified model to explore how primitive nervous systems optimized survival through coordinated movement. Recent advances in computational biology and mathematical modeling are refining our understanding of these coordination mechanisms in simple organisms. The properties of nervous systems using nonequilibrium thermodynamics are proposed in [19, 20]. Economic models have also been proposed to understand which distribution of neurons and connections between them is the most beneficial for the animal’s survival [21].

Model of receptor, activatior, and inhibitor is usually used for chemotaxis analysis, even accounting cellular memory [22]. The directional movement of cells, even in the absence of a nervous system, such as in Trichoplax adhaerens, is facilitated by the elastic forces between them [23]. The size of a organism is limited if it moves in a flow and is unable to choose a direction [24]. However, the nervous system allows the mobile heterotroph to grow to a significantly larger size. At the same time, for small animals, apparently, the nervous system remains unprofitable. This may be one of the reasons why the smallest multicellular organisms like Trichoplax adhaerens or aggregates of cells do not have nervous systems, although the cells are able to exchange signals. We hypothesize that early nervous systems were essential for synchronizing flagellar movements [25] in response to signals from surface sensor cells, allowing multicellular organisms to focus on effective strategies. This raises the question of how cells coordinate their flagellar movements without a central nervous system. For example, one cell can rotate its flagellum clockwise while stimulating another cell to rotate counterclockwise. Bacteria navigate chemical gradients using temporal sampling, while eukaryotic cells sense spatial differences across their body size, crucial for survival. Proteins like Ras, GAF, and GEP regulate ion channels, contributing to movement coordination; in [26] it is shown that activated Ras concentrations are highly dependent on external cAMP gradient around the cell. Understanding these mechanisms sheds light on the evolution of nervous systems in early marine organisms. Moreover, many proteins responsible for the exchange of messages between individual bacteria are similar to chemoreception proteins [27]. Various mating factors of ciliates have been detected including beta-endorphin using by modern mammalians, and at least one of ciliate chemotactic membrane proteins closely resembles opiate receptors of multicellular organisms [28]. Cell-cell recognition, which is necessary, for example, for the fusion of eukaryotic gametes, may also be one of the first innovations in eukaryogenesis associated with cell interactions within the same organism [29].

Our study presents a computational model of a primitive marine organism as a network of motor units linked by elastic springs. We used physiologically relevant time functions for neuronal excitation to simulate behavior and identified key parameters that optimize survival by efficiently locating phytoplankton in competitive environments. With this model, we aim at going beyond our previously proposed models of early nervous systems [30–34] to add more realistic physical behavior. On the other hand, the proposed model extends the existing set of models that can be used to model simple nervous systems at a computationally low cost[35–37]. Our findings suggest that early nervous systems may have enhanced flagellar synchronization and chemotactic coordination among multicellular heterotrophs. We also consider simpler heterotrophic organisms, like choanocytes, as potential precursors to nervous systems, which may have begun forming primitive neural networks. The motor units in our model can mutually excite each other for coordinated movement, with the cytoskeleton serving a role similar to muscle fibers. This study aims to bridge the gap in understanding how early nervous systems evolved to synchronize movements, enhancing survival by optimizing food capture and avoiding harmful stimuli. Through computational simulations, we investigate how these systems may have maximized the survival of simple heterotrophs, offering insights into the emergence of the first nervous systems.

## MODEL

The assumptions on which the model is built:

1. The intercellular connective proteins in the first prototypes of nervous systems were similar to the chemotaxic receptors [38, 39].
2. The first prototypes of nervous systems were needed to synchronize the movements of motor units, such as flagella, cilia and protrusions, in cellular aggregates. This synchronization gave an advantage in the search for food, because otherwise different motor units of the unit could create multidirectional forces, which made it impossible to choose a direction in weak gradients. If the total force was close to zero, then the predator was simply carried away by the flow in the direction where there might not be enough food.
3. As the concentration of substances outside the body sets boundary conditions on receptors, cell movements are determined by their *phases*, which cells try to synchronize by exchanging ions or organic molecules through intercellular contacts. That is, the receptors determine an *effective field* in which each cell chooses a direction, similar to how electric or magnetic dipoles line up along an electric or magnetic field.

The model is described by six groups of equations, describing reception, intracellular polarization, intercellular synchronization, bacterial mats population dynamics, predator mechanics, and energy balance. The formalism of classical mechanics is used.

### I Reception

A boundary condition is set on the animal surface to fix membrane potential depending on local chemoattractant concentration:

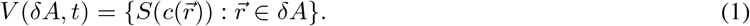

The filling is determined by the classical Langmuir adsorption equation of chemoattractant on the receptors:

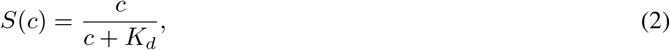

where *c* is the chemoattractant concentration around the receptor, and *K*_*d*_ is the dissociation constant of the complex. In the continuous limit, the concentration is defined for any point of the surface, but actually chemoattractant molecules are discretely distributed over the surface and we used grid approximation described in section IV. At low food concentrations near the heterotroph, sensitivity becomes especially important, and the Langmuir equation approaches a linear dependence. However, at high concentrations, the signals from receptors at different points on the organism body are almost indistinguishable, but the animal survival is not threatened:

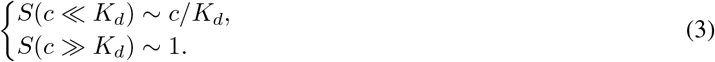

We assume *c* is linearly proportional to the bacterial mat concentration, introduced in equations group IV.

### II Intracellular polarization

Levchenko’s equations from systems biology are widely used to describe distribution of signal pathways activators and inhibitors over cytosol and cellular membrane [40]. This model was developed to calculate chemoattractant polarization vectors driving cellular movement into the steepest gradient rise direction. We use approximation on a polygonal graph (Fig. 1B), using one dimensional results [14] from Levchenko chemoreception-activation-reaction-inhibition pathway kinetics:

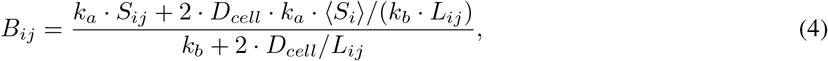

where 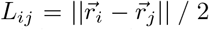 and *k*_*b*_ are activation and inhibition rate constants, *D*_*cell*_ is the diffusion coefficient of the inhibitor in the cells and may significantly differ from bacterial mat diffusion coefficient *D, B*_*ij*_ is inhibitor concentration of *j*-th cell near the channel connecting in with the *i*-th cell. 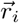 and 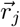 are the coordinate vectors of the adjacent cells. We emphasize that this discrete approximation is rude and shouldn’t be used for processing of experimental intercellular interactions, however, we expect to save qualitative properties of the Levchenko model. Moreover, the number of channels with adjacent cells is much more than one thus the single-channel model is an averaging over all ligand-dependent channels between adjacent cells. *S*_*i*_ and *S*_*ij*_ are defined in the next section.

**Figure 1:**
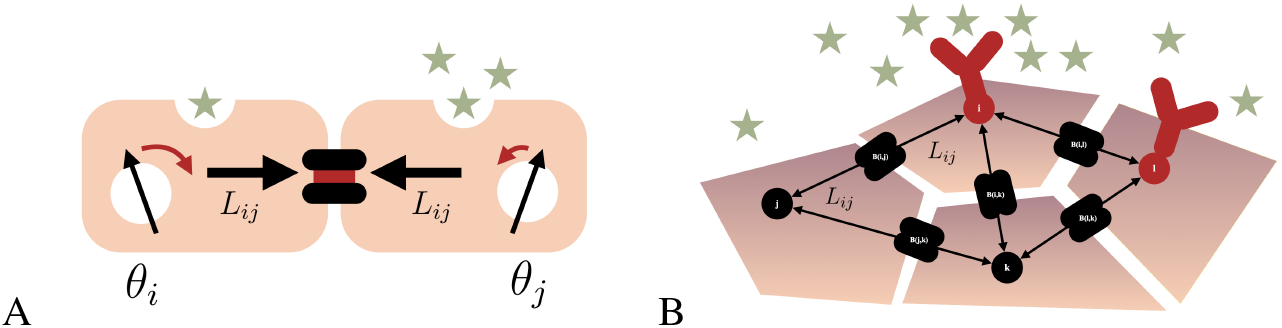
(A) Synchronization of two adjacent cells. (B) Polygon model of receptors, cytoskeletons, and channels.

### III Intercellular synchronization

Synchronization equations of Kuramoto oscillators [41] were modified to account the vectors of target polarization, but the tendency of oscillators to align is conserved. At first glance, it may seem impossible to describe a living organism with a small number of parameters, given the heterogeneity of life. However, many models have been built, such as Lotka-Volterra systems of differential equations, that correspond to experiments. Various distributions from probability theory, Landau oscillator chains, theoretical mechanics or even statistical physics could be applied to the problem of living cells synchronization, but we want to propose a set of simple equations in this section, in future, they should be developed into more precise and biochemically relevant. Excitation of a sensory neuron depends on the distance from the bacterial mat. A variational problem approach is not applicable since the initial and final states of the organism are unknown. However other approaches are available like lifespan maximization or optimization of the size of the motor unit. The phase *θ*_*i*_ of the *i*-th cell is proportional to the rotation speed of the flagellum of the *i*-th effector cell. It may correspond, for example, to the concentration of (de)phosphorylated proteins, whose binding to the basal body increases the number of rotations clockwise or counterclockwise [42] per unit of time. Changes in inhibitor concentration across the entire cell lead to a temporary change in its membrane potential. Expected ratio between the costs of movement and signal transmission. Viscosity allows estimating the steady speed under a constant chemoattractant gradient and is similar to Langevin equations. The direction of the animal movement is ultimately determined by bacterial mats diffusion.

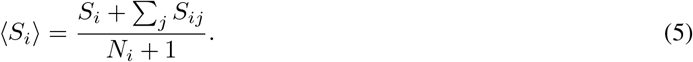

Threshold opening of a channel assumed to occur when *V*_*j*_ *> V*_*i*_ and durates *α*_*dec*_ *· t*_*slow*_:

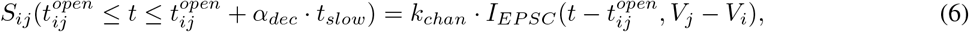

where *k*_*chan*_ is a coefficient to compare chemotactic and intercellular receptors contribution to cellular polarization and 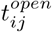 is the time of opening the channel passing the polarization inhibitor from the *j*-th cell to the *i*-th one. Although the model doesn’t assume to contain mature synapses, the general equation for excitatory postsynaptic potential, used, for example, for the jellyfish neuromuscular modeling [16]:

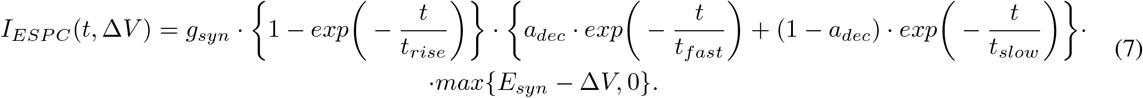

The signal polarization of a cell is the sum of signals from the chemotactic membrane receptor and the channels shared with adjacent cells:

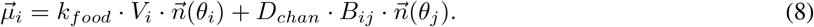

Kuramoto-like update of a phase angle of a cell:

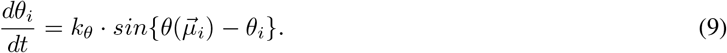

To avoid overload by large numbers, after each iteration phase angle of a cell state is reduced to the equivalent in the limited range:

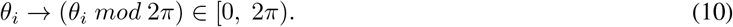

### IV Migration, production, and uptake of bacteria

Common diffusion-advection-reaction equations were used. They are provided, for example, in [43].

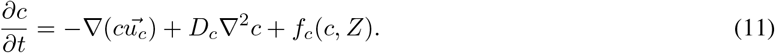

Wavefunction correction multiplier was added to the general *f*_*c*_(*c, Z*) production rule to generate dynamical environment for the predator. Harmonic wave may be the first plausible approximation for periodic dynamics of the bacterial mats:

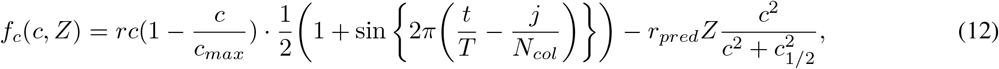

where *r* is bacterial mat production rate constant, *c*_*max*_ is the concentration of saturation, *T* is the time period of the wave and *N*_*col*_ is the number of grid columns in a simulation. *r*_*pred*_ corresponds to uptake of food by the heterotroph and *Z* is taken equal to number of animal cells covering the grid cell at the moment. The initial bacterial mat distribution was generated randomly: in each cell, the initial concentration was randomly selected from 0 to *c*_*max*_, because diffusion quickly organized the initial chaos, but it generated a variety of patterns, allowing qualitatively different predator trajectories to be discovered.

### V Predator mechanics

Of course, these motor units cannot be considered as cells that are connected by springs — the structure should be considered as an averaging, which can have at least three interpretations: (1) a motor unit is actually the averaging of a group of cells, and the fibers connecting the motor units are selected to average the elastic forces between, for example, the centers of masses of these groups of cells; (2) a motor unit is a group of cells, a fiber is a giant axon connecting two groups of nerve cells [44]; (3) a motor unit is the center of a single cell, and the fiber is a fragment of the cytoskeleton directed from this center to the membrane channel, after which another cell begins. The signed rotation speed of the flagellum of a surface effector cell assumed to be proportional to the projection of cell phase vector on the axis connecting the animal center of masses with the cell. The force exerted by the rotation of the flagellum of a surface cell:

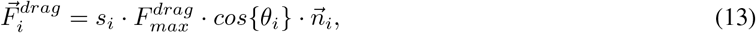

where *s*_*i*_ ∈ *{* 0, 1*}* is the indicator of whether a cell is sensory, 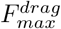 is the maximal drag force from flagellar rotation, and 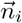 is the flagellum unit direction outward vector. Springs (s — spring) between neurons create forces:

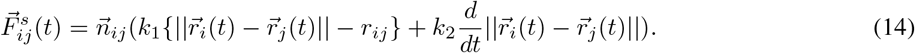

The total elastic force acting on the *i*-th cell:

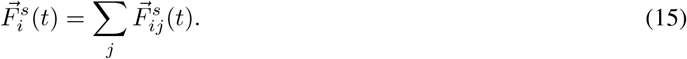

Viscosity force in general:

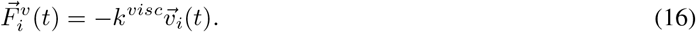

The total force acting on an individual neuron produces its acceleraton:

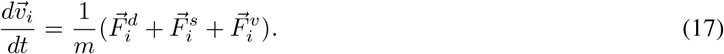

In water for a bacterium, *Re* ∼ 3 *·*10^−5^, so for a small multicellular organism, *Re* ≪ 1. In the case of constant chemoattractant gradient surrounding the animal, its established velocity is [24]

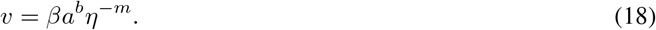

A more accurate description of the movement of the cilia, taking into account the frequency of push and return, can be taken, for example, from [45], and with respect to multicilia interaction from [46].

### VI Energy balance of a heterotroph

The cost of motor work and the energetics of the flagellum are connected to the thrust force and costs for cilia movement [47]:

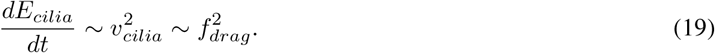

The spread of the gradient of signaling particles — metal ions or organic neurotransmitters, ATP — can occur through diffusion without energy costs or through active transport requiring energy. The survival of the animal is determined by an energy function *W*, which increases from consuming food particles and is spent on the work of “muscles” with a multiplier and the “nervous system”. The differential term corresponds to the work of creating a Ca^2+^ concentration gradient, and the integral term to the proportional contraction of “muscles” and transmission of the activation potential. It is assumed that one unit of consumed food provides the heterotroph with energy, and over one unit of time, the predator consumes a mass of food *p*(*t*), while homeostasis, the work of the sodium-potassium pump, etc., are required at a cost h (homeostasis) per unit of time: (?) At the beginning of the simulation, an initial condition *W* (0) is set. The organism will die as soon as *W* (*t*) *<* 0. The ratio *ϕ ·* (*ω*^−1^ + *λ*^−1^) can be called the heterotroph efficiency coefficient. The ratio *ϕ* : *γ* : *ω* can be estimated from [24, 29].

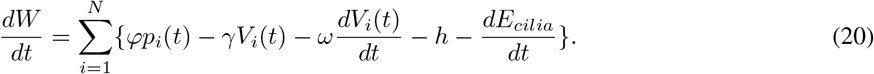

For comparison, without a nervous system, a spherical heterotroph spends less energy [24] but could gather less food:

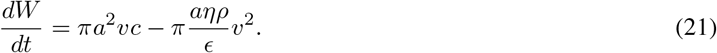

## RESULTS

The longer the organism lives, the higher the probability of reproduction, so the average lifespan of the animal could be selected as the parametere to maximizd (assuming the organisms do not survive to natural death):

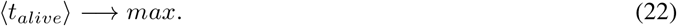

Another approach is to fix simulation time and collect average remained predator energy after each simulation:

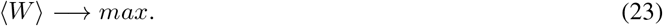

Both approaches, among all the tested activation-inhibition ratios for the fixed remaining parameters of the model, gave the best result at *k*_*a*_*/k*_*b*_ = 0.1 (Fig. 2), after which a sharp decline began. These results show that excessively high excitability is dangerous, while fast synchronization gives more than an abrupt reaction to a locally high concentration of food. So, evolution had to establish the optimal ratio between excitability, relaxation, and synchronization rates. Moreover, it seems to occur the plateu for the little activation rates that emerged due to the finite size of the grid cells used in the simulations.

**Figure 2:**
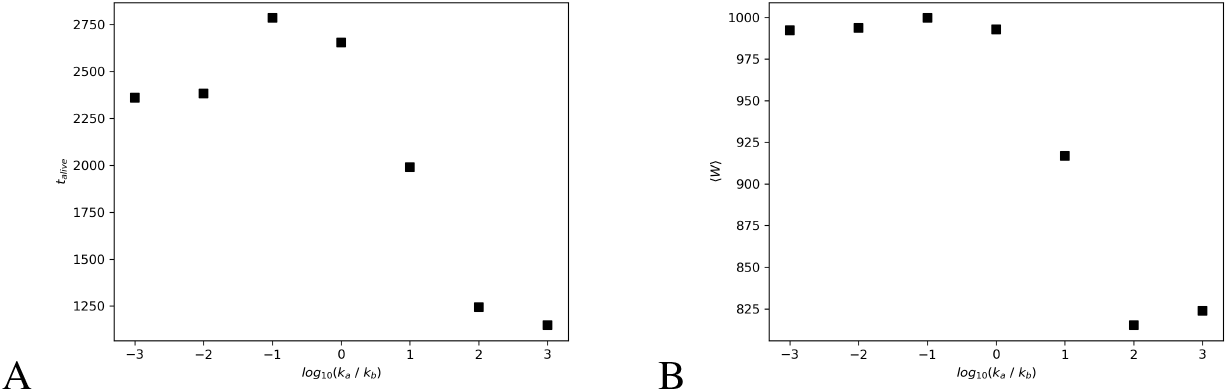
Both types of computations were conducted 3 times. (A) The dependence of the average survival time on the ratio of activation and inhibition rates of membrane receptors. (B) The dependence of the average energy reserve of the organism on the same parameter.

## SOFTWARE

Although the distribution of neuron types in even highly symmetric Cnidarians varies significantly along the axis [48], in each of the simulations we simulated a single symmetrical distribution of neurons around the center, and the predator was assumed to be flat. shows how synchronization between cells increases as activation-inhibition ratio becomes greater. Colors of motor units correspond to *θ*_*i*_ and colors of grid cells correspond to the bacterial mat concentrations.

**Figure 3:**
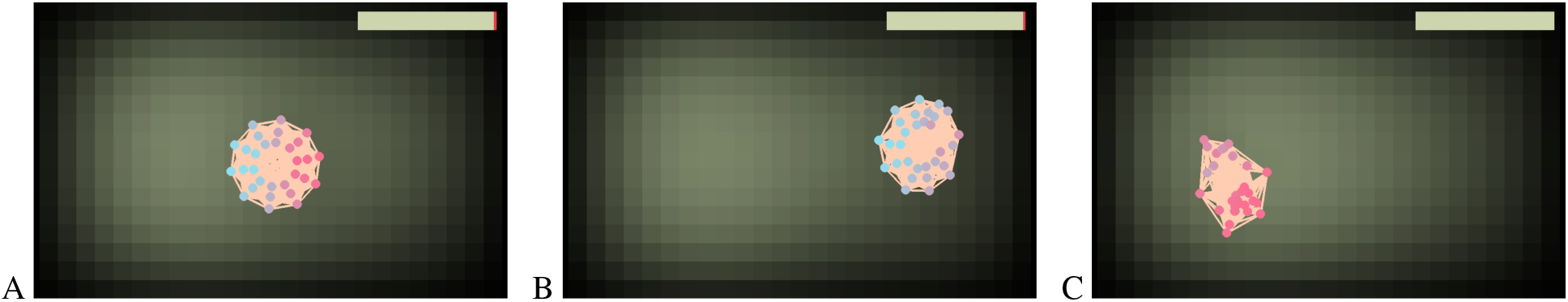
Simulation screens for *k*_*a*_*/k*_*b*_ = (A) 10^−3^; (B) 10^−2^; (C) 10^−1^.

We consider the software as an open platform for studying the evolution of the nervous systems. It is written in Python 3 using Pygame library. Link to the repository can be found there: https://github.com/YegorVasilenko/multicellular_chemotaxis. Anyone can select parameter values for the energy benefits of food, the energetic costs of signal transmission or muscle contraction, and any other model parameters.

## CONCLUSION

Even such a simple model utilizing basics of mechanics and elasticity, crude approximations of activation-inhibition chemical kinetics, and schematic representation of ion channels generates complex behavior. The lattice model of bacterial mat limits the realism of the model, but if one will reduce grid step, the simulation should become more plausible. It is interesting how an increase in the activation constant compared to the inhibition constant first increases the survival rate of a predator, and then decreases it. Also, as this ratio increases, the acceleration of cell synchronization is noticeable, which, however, eventually leads to too sharp “throws” of the animal to unexplored and, possibly, critically low concentrations of food.

The presented algorithm and similar ones for controlling multicellular predators can be used outside of biology, for example, to create scalable and flexible multi-module robots that can be used to collect toxins or transport cargo. Although the model is a rude approximation and ignores many processes underlying both motion and signal transduction within multicellular organisms, we suggest the proposed approach may provide a platform for further development of the six systems of equations.

## REFERENCES

[1] Jeremy E Niven and Lars Chittka. “Evolving understanding of nervous system evolution”. Current Biology 26.20 (2016), R937–R941.

[2] Michael G Paulin and Joseph Cahill-Lane. “Events in early nervous system evolution”. Topics in Cognitive Science 13.1 (2021), pp. 25–44.

[3] Dirk Bucher and Peter AV Anderson. Evolution of the first nervous systems–what can we surmise? 2015.

[4] Graham E Budd. “Early animal evolution and the origins of nervous systems”. Philosophical Transactions of the Royal Society B: Biological Sciences 370.1684 (2015), p. 20150037.

[5] Gáspár Jékely, Jordi Paps, and Claus Nielsen. “The phylogenetic position of ctenophores and the origin (s) of nervous systems”. Evodevo 6 (2015), pp. 1–9.

[6] Fred Keijzer. “Moving and sensing without input and output: early nervous systems and the origins of the animal sensorimotor organization”. Biology & Philosophy 30.3 (2015), pp. 311–331.

[7] Benjamin J Liebeskind et al. “Complex homology and the evolution of nervous systems”. Trends in ecology & evolution 31.2 (2016), pp. 127–135.

[8] Gáspár Jékely. “The chemical brain hypothesis for the origin of nervous systems”. Philosophical Transactions of the Royal Society B 376.1821 (2021), p. 20190761.

[9] Leonid L Moroz. “Multiple origins of neurons from secretory cells”. Frontiers in Cell and Developmental Biology 9 (2021), p. 669087.

[10] Leonid L Moroz and Daria Y Romanova. “Alternative neural systems: what is a neuron?(ctenophores, sponges and placozoans)”. Frontiers in Cell and Developmental Biology 10 (2022), p. 1071961.

[11] Carl Frederick Abel Pantin. “Croonian Lecture-The elementary nervous system”. Proceedings of the Royal Society of London. Series B-Biological Sciences 140.899 (1952), pp. 147–168.

[12] Yoshiki Kuramoto and Yoshiki Kuramoto. Chemical turbulence. Springer, 1984.

[13] Tilmann Glimm and Daniel Gruszka. “Modeling the interplay of oscillatory synchronization and aggregation via cell–cell adhesion”. Nonlinearity 37.3 (2024), p. 035016.

[14] Herbert Levine, David A Kessler, and Wouter-Jan Rappel. “Directional sensing in eukaryotic chemotaxis: a balanced inactivation model”. Proceedings of the National Academy of Sciences 103.26 (2006), pp. 9761–9766.

[15] Luquan Ren et al. “Biology and bioinspiration of soft robotics: Actuation, sensing, and system integration”. Iscience 24.9 (2021).

[16] Fabian Pallasdies et al. “From single neurons to behavior in the jellyfish Aurelia aurita”. Elife 8 (2019), e50084.

[17] Michiya Kamio and Charles D Derby. “Finding food: how marine invertebrates use chemical cues to track and select food”. Natural Product Reports 34.5 (2017), pp. 514–528.

[18] Iva Kelava, Fabian Rentzsch, and Ulrich Technau. “Evolution of eumetazoan nervous systems: insights from cnidarians”. Philosophical Transactions of the Royal Society B: Biological Sciences 370.1684 (2015), p. 20150065.

[19] Beren Millidge, Anil Seth, and Christopher L Buckley. “A mathematical walkthrough and discussion of the free energy principle”. 2108.13343 (2021).

[20] Biswa Sengupta, Martin B Stemmler, and Karl J Friston. “Information and efficiency in the nervous system—a synthesis”. PLoS computational biology 9.7 (2013), e1003157.

[21] Vincenzo Nicosia et al. “Phase transition in the economically modeled growth of a cellular nervous system”. Proceedings of the National Academy of Sciences 110.19 (2013), pp. 7880–7885.

[22] Richa Karmakar et al. “Cellular memory in eukaryotic chemotaxis depends on the background chemoattractant concentration”. Physical Review E 103.1 (2021), p. 012402.

[23] Carolyn L Smith et al. “Coherent directed movement toward food modeled in Trichoplax, a ciliated animal lacking a nervous system”. Proceedings of the National Academy of Sciences 116.18 (2019), pp. 8901–8908.

[24] William W Crockett et al. “Physical constraints during Snowball Earth drive the evolution of multicellularity”. Proceedings of the Royal Society B 291.2025 (2024), p. 20232767.

[25] Li Xie et al. “Bacterial flagellum as a propeller and as a rudder for efficient chemotaxis”. Proceedings of the National Academy of Sciences 108.6 (2011), pp. 2246–2251.

[26] Yougan Cheng, Bryan Felix, and Hans G Othmer. “The roles of signaling in cytoskeletal changes, random movement, direction-sensing and polarization of eukaryotic cells”. Cells 9.6 (2020), p. 1437.

[27] Yuri B Shmukler and Denis A Nikishin. “Non-neuronal transmitter systems in bacteria, non-nervous eukaryotes, and invertebrate embryos”. Biomolecules 12.2 (2022), p. 271.

[28] GO Mackie. “The elementary nervous system revisited”. American Zoologist 30.4 (1990), pp. 907–920.

[29] Kirsty Y Wan and Gáspár Jékely. “Origins of eukaryotic excitability”. Philosophical Transactions of the Royal Society B 376.1820 (2021), p. 20190758.

[30] Anton V Sinitskiy. “Simplest Model of Nervous System. I. Formalism”. bioRxiv (2023). DOI: 10.1101/2023.11.23.568481.

[31] Anton V Sinitskiy. “Simplest Model of Nervous System. II. Evolutionary Optimization”. bioRxiv (2023). DOI: 10.1101/2023.11.24.568590.

[32] Anton Sinitskiy. “Simplest Model of Nervous System. III. Partial Optimization”. bioRxiv (2024). DOI: 10.1101/2024.06.20.599964.

[33] Anton Sinitskiy. “Simplest Model of Nervous System. IV. General Solution”. bioRxiv (2024). DOI: 10.1101/2024.07.10.603010.

[34] Anton V Sinitskiy. “Making Sense of Neural Networks in the Light of Evolutionary Optimization”. bioRxiv (2023). DOI: 10.1101/2023.11.27.568922.

[35] Yuval Tassa et al. “Deepmind control suite”. 1801.00690 (2018).

[36] Greg Brockman et al. “Openai gym”. 1606.01540 (2016).

[37] Samuel Schmidgall, Catherine Schuman, and Maryam Parsa. “Biological connectomes as a representation for the architecture of artificial neural networks”. bioRxiv (2022). DOI: 10.1101/2022.09.30.510374.

[38] Asba Tasneem et al. “Identification of the prokaryotic ligand-gated ion channels and their implications for the mechanisms and origins of animal Cys-loop ion channels”. Genome biology 6 (2005), pp. 1–12.

[39] Ákos Nemecz et al. “Emerging molecular mechanisms of signal transduction in pentameric ligand-gated ion channels”. Neuron 90.3 (2016), pp. 452–470.

[40] Herbert Levine and Wouter-Jan Rappel. “The physics of eukaryotic chemotaxis”. Physics today 66.2 (2013), pp. 24–30.

[41] Forest O Mannan, Miika Jarvela, and Karin Leiderman. “Minimal model of the hydrodynamical coupling of flagella on a spherical body with application to Volvox”. Physical Review E 102.3 (2020), p. 033114.

[42] Birgit E Scharf et al. “Control of direction of flagellar rotation in bacterial chemotaxis”. Proceedings of the National Academy of Sciences 95.1 (1998), pp. 201–206.

[43] Jonathan Reid Woodward, Jonathan William Pitchford, and Martin Alan Bees. “Physical flow effects can dictate plankton population dynamics”. Journal of the Royal Society Interface 16.157 (2019), p. 20190247.

[44] Nadia Riebli and Heinrich Reichert. “The first nervous system”. The Wiley Handbook of Evolutionary Neuroscience (2016), pp. 125–152.

[45] John R Blake and Michael A Sleigh. “Mechanics of ciliary locomotion”. Biological Reviews 49.1 (1974), pp. 85–125.

[46] Shay Gueron and Nadav Liron. “Ciliary motion modeling, and dynamic multicilia interactions”. Biophysical journal 63.4 (1992), pp. 1045–1058.

[47] Shay Gueron and Konstantin Levit-Gurevich. “Energetic considerations of ciliary beating and the advantage of metachronal coordination”. Proceedings of the National Academy of Sciences 96.22 (1999), pp. 12240–12245.

[48] Hiroshi Watanabe, Toshitaka Fujisawa, and Thomas W Holstein. “Cnidarians and the evolutionary origin of the nervous system”. Development, growth & differentiation 51.3 (2009), pp. 167–183.

